# Design and Fabrication of Petri Dish Optimized for High-Frequency Acoustic Microscopy Using 3D Printing

**DOI:** 10.64898/2025.12.12.693677

**Authors:** Tuna Pesen, Beyza Ceren Eren, Bora Akgun

## Abstract

High-frequency imaging in acoustic microscopy requires the use of petri dishes with thin and acoustically transparent bottoms to minimize signal attenuation and distortion. Commercially available options suitable for this purpose are often expensive or limited in compatibility with custom setups. Here, we present a cost-effective and easily customizable alternative developed in our laboratory, consisting of a 3D-printed frame combined with commercially available stretch film. The stretch film serves as an ultrathin (8*µm* thickness) base that supports efficient high-frequency acoustic transmission, enabling clear and reliable imaging. Beyond acoustic microscopy, the adaptable design is also compatible with other modalities such as optical tweezers and fluorescence microscopy. This versatility significantly enhances the value of the proposed design, offering a practical solution for diverse experimental needs.

## Introduction

Acoustic microscopy is an advanced imaging technique that employs high-frequency sound waves to probe the structural, mechanical, and elastic properties of biological samples at microscale resolution.^1,2^ Unlike optical or electron microscopy, which rely on electromagnetic interactions, acoustic microscopy utilizes the propagation of acoustic waves, typically in the MHz to GHz range, offering unique contrast mechanisms based on material density and stiffness.^3^ This enables non-invasive, label-free imaging of soft and heterogeneous biological specimens, such as cells, tissues, and extracellular matrices.^4^

In biophysical research, acoustic microscopy allows for the quantification of parameters such as acoustic impedance, sound velocity, and attenuation, which are directly related to the elastic modulus and viscoelastic behavior of the specimen. These measurements are particularly valuable in studying processes like cell differentiation, disease progression, tissue remodeling, and the external effects such as heat, drugs, or pathological diagnosis.^5–10^

Recent developments in scanning acoustic microscopy (SAM) and high-frequency transducers have further enhanced the spatial resolution and sensitivity of the technique, making it possible to investigate subcellular features and dynamic changes in cell mechanics.^11^ Given its ability to interrogate mechanical properties and minimal sample preparation (< 1 hour), acoustic microscopy continues to expand its role in mechanobiology, pathology, and biomedical engineering, providing critical insights into the physical principles that govern cellular structure and function.^12–14^

A critical consideration in acoustic microscopy, especially when working at ultra-high frequencies (e.g., >100 MHz), is the acoustic coupling and transmission efficiency between the transducer and the biological sample.^15^ To minimize signal attenuation and phase distortion, the sample must be mounted on a petri dish or substrate with an extremely thin bottom layer, often made of acoustically transparent materials such as polystyrene or polymer films.^16^ These thin-bottom dishes allow for efficient propagation of high-frequency acoustic waves, preserving both axial and lateral resolution, and are essential for maximizing image quality and measurement accuracy in cellular-level investigations.^15,16^

Despite the availability of commercial petri dishes with thin, acoustically transparent bottoms, their high cost and lack of compatibility with custom-designed experimental systems pose significant limitations for research laboratories. In this study, we present a low-cost and easily customizable solution developed in our laboratory, consisting of a 3D-printed frame and commercially available stretch film. The design provides a reliable and effective alternative that supports efficient acoustic wave transmission while allowing flexible design modifications tailored to specific experimental needs. In addition, the flexible design ensures compatibility with multiple measurement modalities beyond acoustic microscopy, including optical tweezers and fluorescence microscopy. This versatility further enhances the value of the custom-made design for advanced research applications.

### Theory of Acoustic Impedance

Acoustic impedance (Z) is defined as the resistance a material offers to the propagation of sound waves. The definition is:

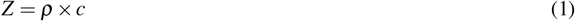

where *ρ* is the density of the medium and *c* is the speed of propagation of sound in that medium. When the density or speed of sound in an environment changes, the acoustic impedance also changes. Acoustic impedance modeling allows for analysis of the acoustic behavior of a sample or multilayer structure at different frequencies.^17,18^ Wave reflection and transition analyses form the basis of this mode. As the impedance difference between the two media increases, the amount of energy reflected decreases. This situation is expressed as follows:

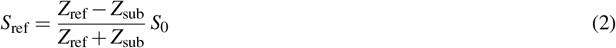

Here, the formula involving *S*_ref_, calculated using the acoustic impedance of water, *Z*_ref_, and that of the substrate, *Z*_sub_, provides the reflection ratio at the interface and, *S*_0_ is the transmitted signal from the probe. The direction of the signals can be seen in Fig.1. The *reference* measurement is essential for the initial calibration of the system. Generally, purified water or ultrasound gel is used as the *reference* signal. In acoustic imaging, these strong reflections appear in images as bright or high-signal regions. When a sound wave reaches the interface (boundary) of two media with different acoustic impedance, part of the wave is reflected (returned), and part passes to the other medium (transmitted). The ratio of this reflection and transmission is directly affected by the magnitude of the acoustic impedance difference that separates the two media.

**Figure 1.**
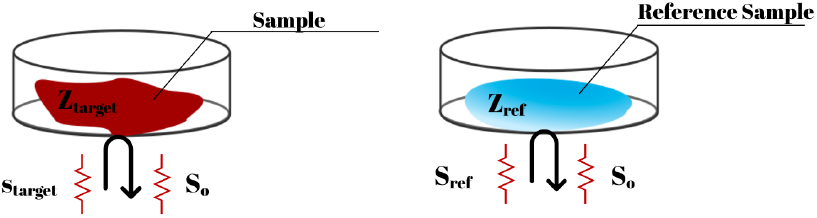
Transmitted and the reflected acoustic signals from a sample.

The substrate acts as the basic surface on which the structure or cell samples to be tested are placed. The substrates used in acoustic systems not only serve as physical carriers; they also directly affect the behavior of acoustic waves, such as propagation, reflection, and refraction. Hard substrates with high impedance (e.g. glass, sapphire) allow the reflection of high-frequency waves,^19,20^ while polymer substrates with low impedance (e.g. PDMS) can offer smoother transition profiles and better compatibility with biological samples.^21,22^ The signal reflected from the target tissue (*Z*_target_), *S*_target_ can be calculated as follows:

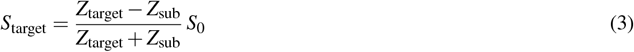

This equation tells us how the tissue’s own acoustic impedance creates an obstacle relative to the substrate impedance. Here, the larger the difference between the *Z*_target_ and the *Z*_sub_, the more severe the *S*_target_ signal will be the interval. The impedance of target tissue (*Z*_target_) can be calculated using the intensity of the reflected signal from target (*S*_target_) and the known substrate impedance (*Z*_sub_) as follows:

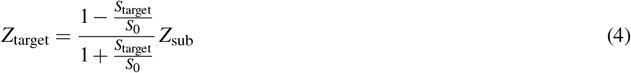

This calculation enables the quantitative determination of acoustic impedance of tissue or material to understand their mechanical features.

### Methods: Petri Dish Design and Production

In this study, a low-cost and customizable petri dish with a thin and acoustically transparent bottom was designed and manufactured for use in high-frequency acoustic microscopy experiments. The design consists of three main parts: a base frame, a compression ring that secures the stretch film, and a lid. The base frame ensures that the film surface remains fixed and flat, while the compression ring mechanically stretches and holds the film in place. The lid is used to prevent evaporation and protect the sample during experiments. Figure 2 shows the technical drawings of each individual part.

**Figure 2.**
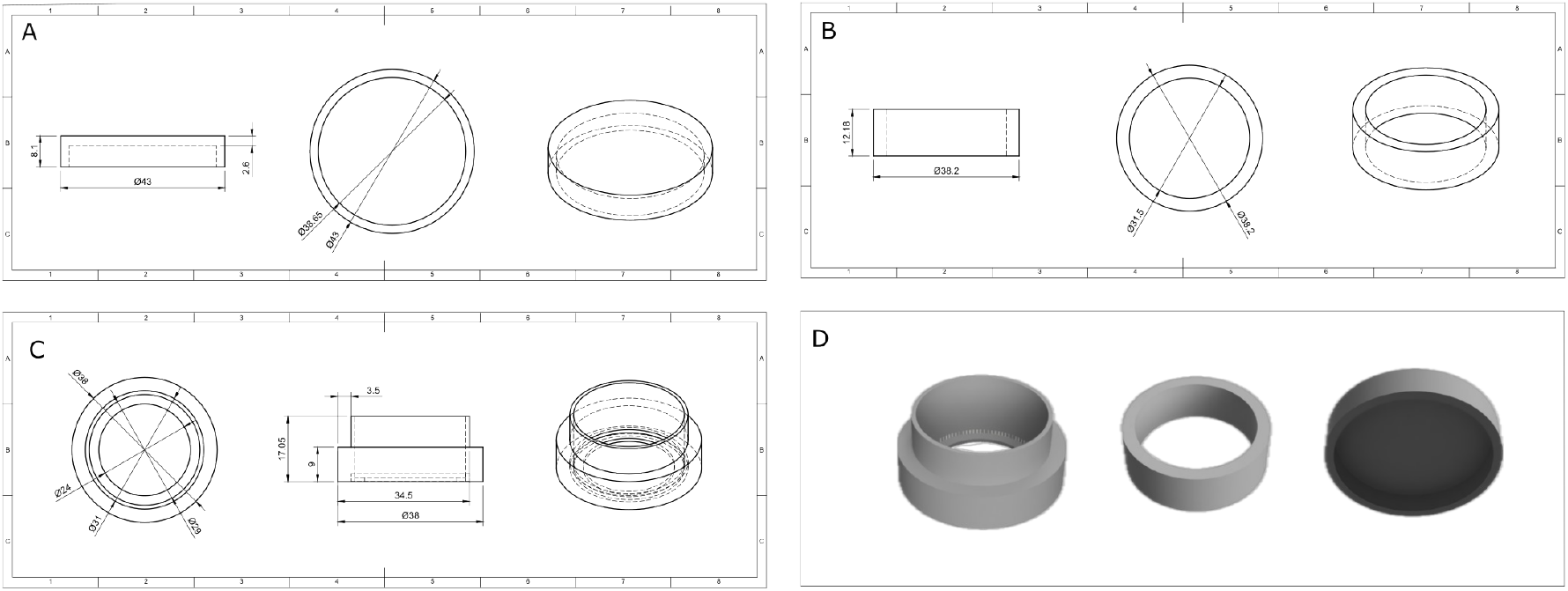
Technical drawings of a) the lid, b) the compression ring, c) the body and, d) renders of parts of the petri dish.

All parts were modeled using the [Autodesk Fusion 360](https://www.autodesk.com/products/fusion-360/overview) software and printed using the Monoprice Maker Select V2 3D printer with Fused Deposition Modeling (FDM) technology (see Fig.3). The slicing process was carried out using [Ultimaker Cura](https://ultimaker.com/software/ultimaker-cura/) software, where appropriate printing settings were configured, and the corresponding G-code was generated for 3D printing. Figure 4 presents the complete rendered assembly of all the petri components.

**Figure 3.**
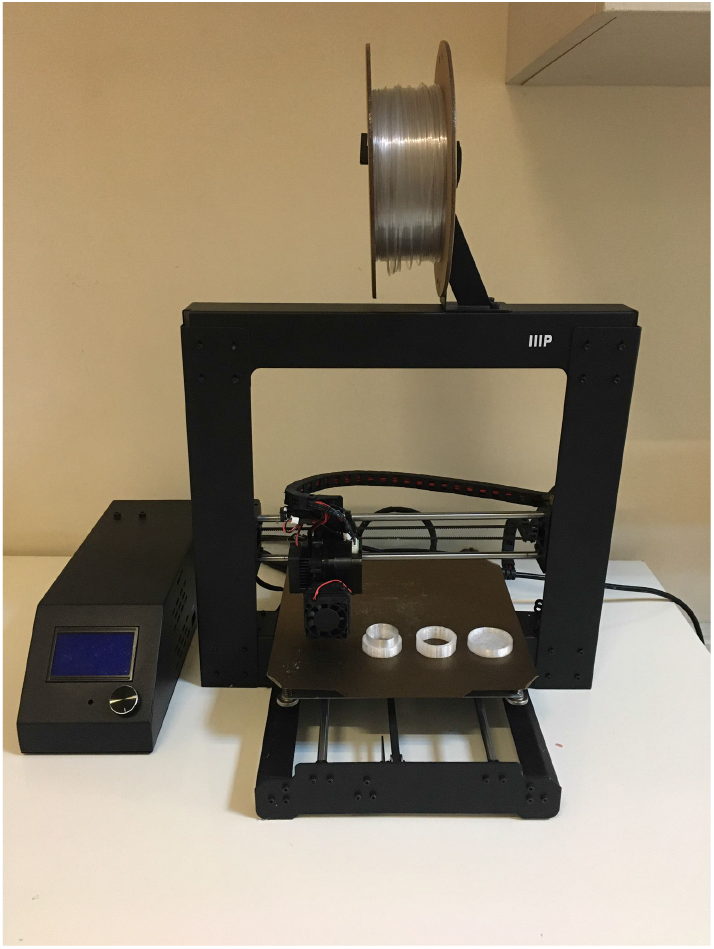
Monoprice Maker Select V 3D printer.

**Figure 4.**
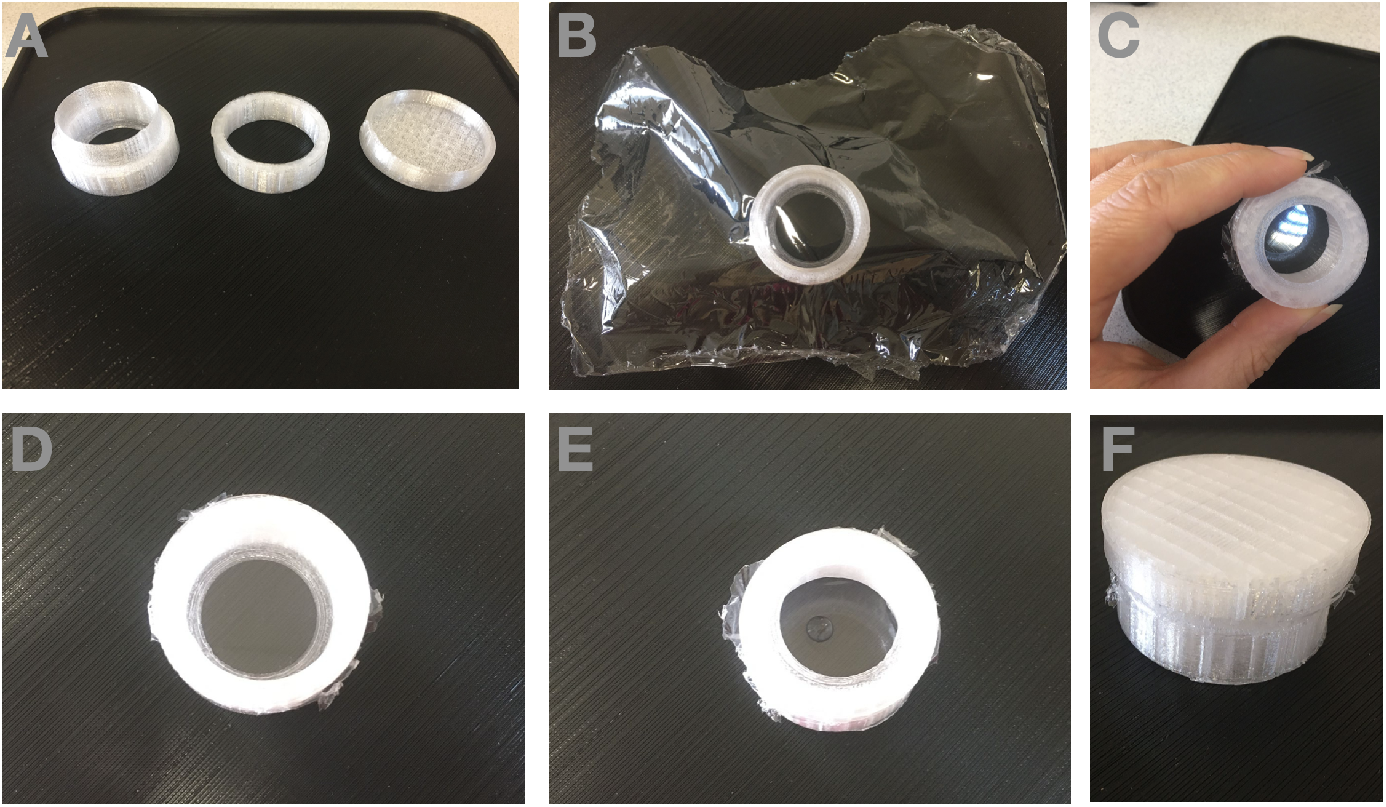
Assembly of the petri dish from a-c) the printed bodies and the stretch film, d) assembled petri dish, e,f) water droplet on the petri as a test sample.

A translucent Hyper PETG filament, known for its great mechanical strength and semitransparency, was chosen for printing. The printing parameters were as follows: 0.3 mm diameter nozzle, 0.15 mm layer height, 240^*°*^C printing temperature, 60^*°*^C build plate temperature, 20/100 infill density and 70 mm/s printing speed. In ultra-quick mode, it took about 40 minutes to print a whole petri dish set and, also 65 minutes in standard mode.

The dimensions of the printed petri dish is 38.0 mm in outer diameter, 31.0 mm in inner diameter, and 17.0 mm in height. The compression ring (clamping piece) has an outer diameter of 38.2 mm, an inner diameter of 31.5 mm, and a height of 10.1 mm. The lid measures 43.0 mm in outer diameter, 38.6 mm in inner diameter, and 8.1 mm in height. The design includes these three main components, all sized accordingly. The usable volume of the petri dish, calculated based on its internal diameter (38.0 mm) and height (17.0 mm), is approximately 19.3 mL.

The total filament used to print all components was approximately 13 grams. Considering the market cost of Hyper PETG filament as $20.1 per kilogram, the material cost per petri dish was calculated to be around $0.25. This cost-effectiveness allows for broader accessibility and rapid prototyping in laboratory environments, especially where budget constraints are a concern.

The stretch film was manually stretched over the base frame and secured with the compression ring. The film surface was ensured to be smooth, taut, and wrinkle-free. The resulting assembly provides an inexpensive and easily manufacturable alternative petri dish suitable for the transmission of high-frequency acoustic signals.

## Measurements and Results

We successfully fabricated thin-bottom petri dishes using a custom-designed 3D-printed-frame and commercially available 8-micron-thick stretch film ([COOK EDT Strec Film PVC Sari 8 Mic](https://magaza.cook.com.tr/cook-edt-strec-film-pvc-sari-8-mic-plastik-testereli)). The resulting dishes provided an acoustically transparent and mechanically stable platform suitable for high-frequency imaging applications. The structural integrity showed that the stretch film, when properly tensioned on the 3D printed frame, maintained its consistent tension. To evaluate acoustic performance, we performed high-frequency imaging using a scanning acoustic microscope operating at frequencies 80 MHz and 320 MHz. Our custom-made dishes demonstrated excellent acoustic transmission, allowing clear visualization of test samples with minimal signal distortion or attenuation. We measured blood smear, as a test sample, in our custom-made petri dish using 320 MHz transducer as given in Fig.5. To test custom-made petri in acoustic imaging using an 80 MHz transducer, Figure 7 demonstrates the measurement of the targetted area of the coin and the corresponding acoustic signal intensity and impedance maps. The intensity and acoustic impedance images gathered using commercial petri (Fig.7c,d) and custom-made petri (Fig.7e,f) gave almost the same image quality to eye, however, since the thickness differences, the acoustic impedance of the background slightly differs, as can be seen on the Fig.7d and f according to the colorbar.

**Figure 5.**
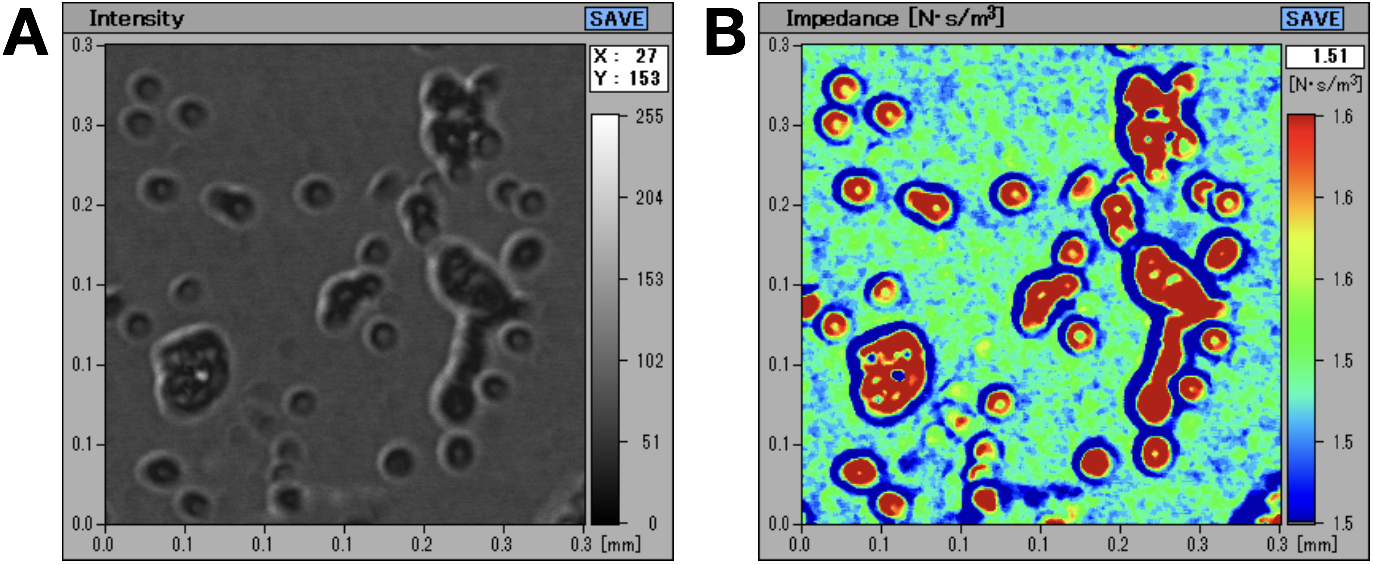
Red blood cell smear measured with 320 MHz transducer on a custom-made petri with 8*µ*m thin bottom. we measured a coin embedding it in an ultrasound gel as shown in Fig. 6.

**Figure 6.**
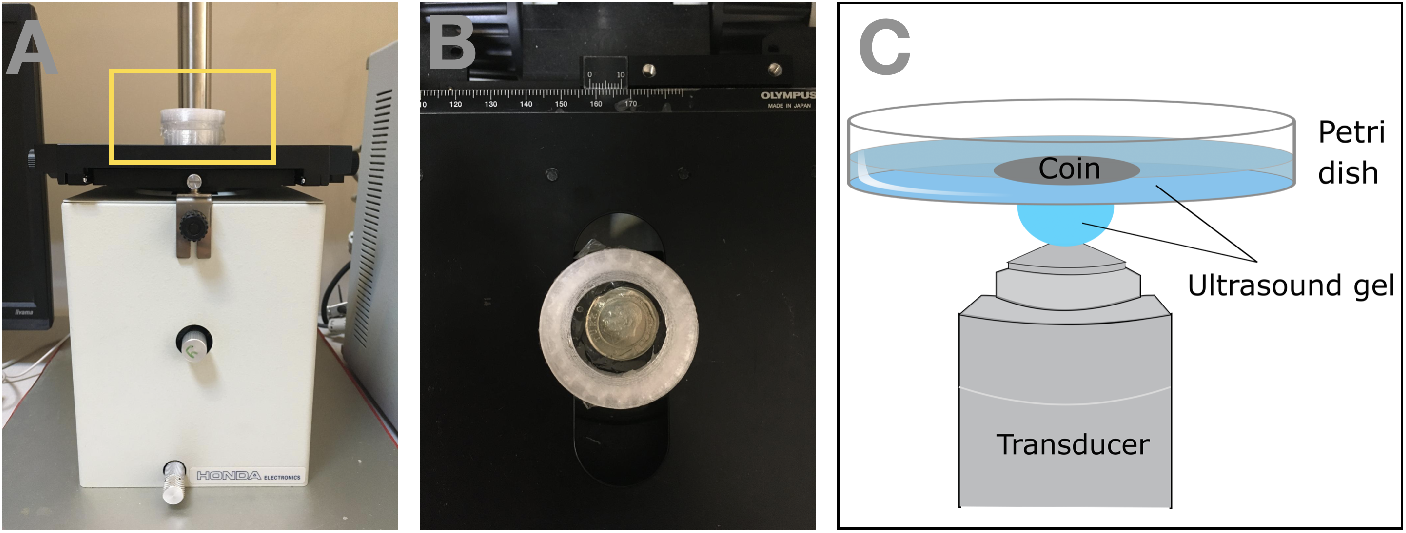
Measurement of a coin on a) the SAM system using b) our custom-made petri dish. The coin was measured by being embedded into ultrasound gel.

**Figure 7.**
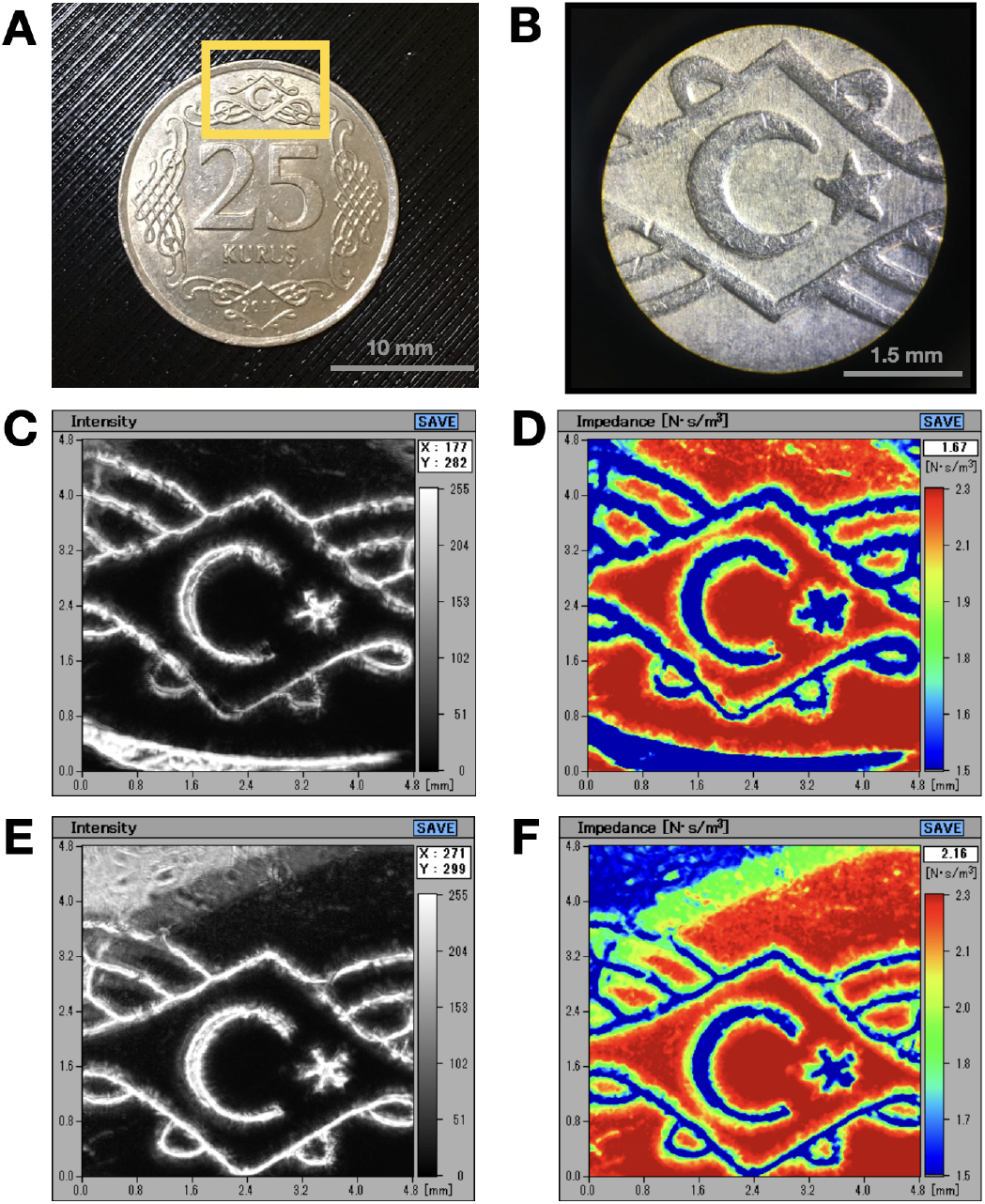
a) The image of the coin an the targeted part shown in yellow square, b) The bright field image of the targetted part, c) Acoustic signal intensity map and d) acoustic impedance map of the coin scanned with 80 MHz transducer on a custom-made petri dish with 8 *µ*m thin bottom and e,f) the images gathered using a commercial petri dish with 50 *µ*m thin bottom.

Optical transmittivity of the bottom of the petri dishes was tested using light microscopy. Red blood cell (RBC) sample in phosphate buffered saline (PBS) was viewed (see Fig.8a) using 100X objective EC “Plan-Neofluar” 100x/1.3 Oil Iris M27 with a numerical aperture of 1.3 and transmittance of about 50% at 1064 nm laser wavelength.. We also tested the usage of the petri for optical tweezers experiments. To do this, Zeiss PALM Micro Tweezers system was used. It has 1064 nm wavelength laser of and the laser power of 800 mW behind the objective. Optical tweezers enable to manipulate micron-sized objects with laser.^23–25^ In Fig.8b, an optically trapped white blood cell was shown. Additionally, the design allows exploration of drug response, as it has a lid that enables the addition of substances during experiments. For example, in Fig.9, we showed the effect of Aspirin (Bayer, 500 mg tablet) on red blood cells by turning them into RBC ghosts with time.

**Figure 8.**
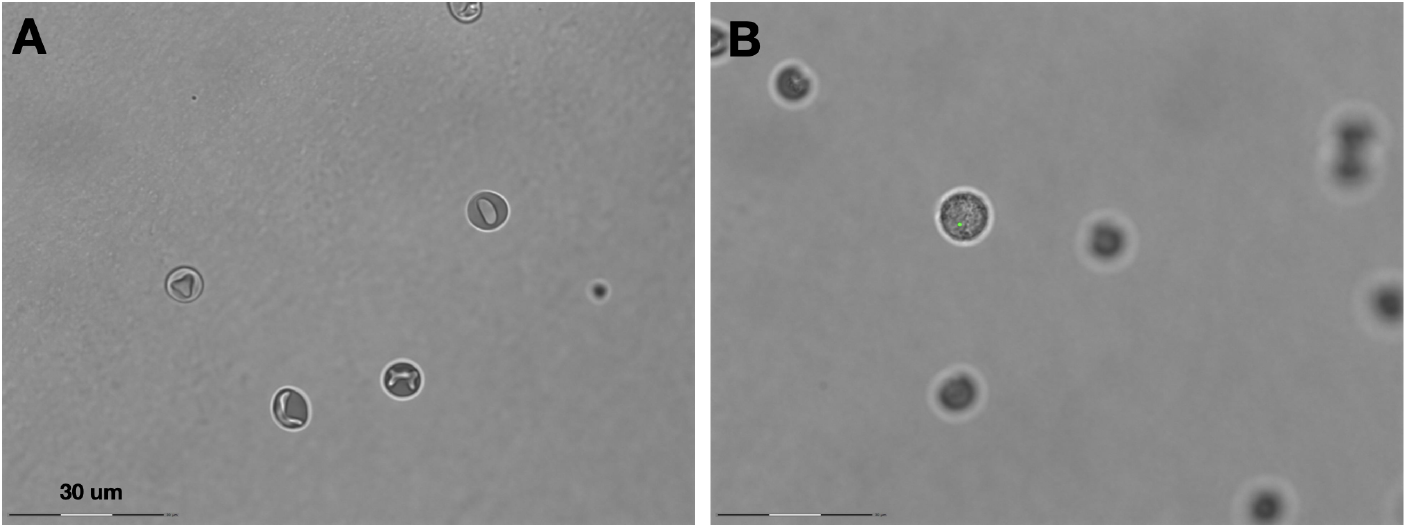
Image of a) red blood cells and b) optically trapped white blood cell on the 8 *µm*-thick petri dish, using a bright field microscopy. The green dot indicates the location of the laser trap. The scale bar is 30 *µm*.

**Figure 9.**
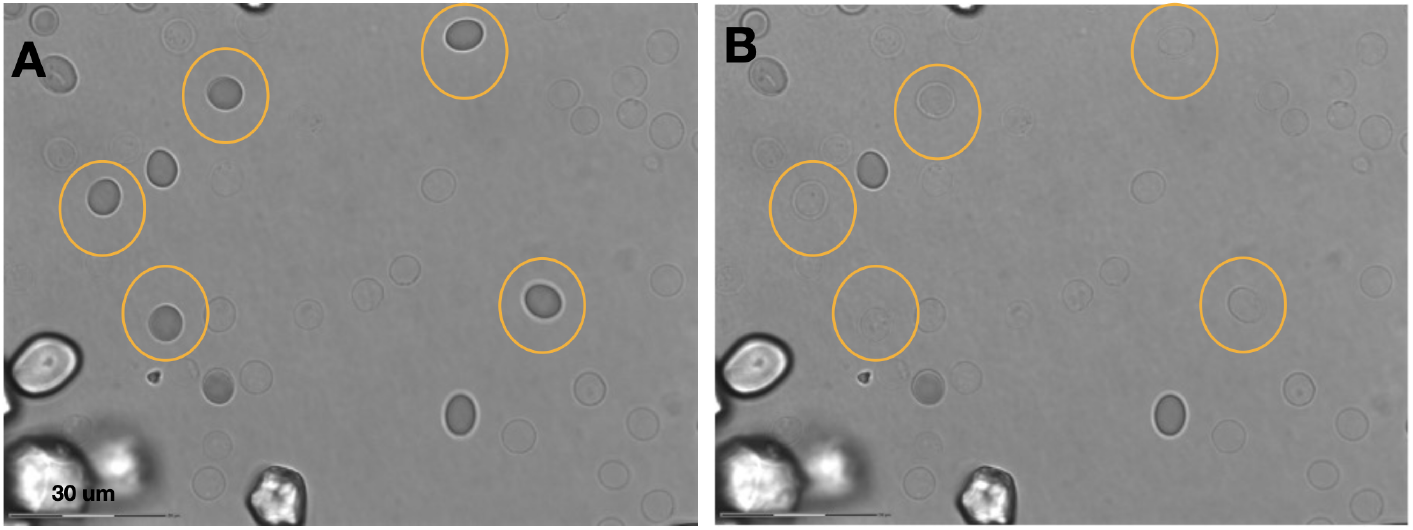
a) RBCs after 35 min Aspirin was added, b) RBCs after 55 min Aspirin was added. The time interval between (a) and (b) was 20 minutes. In that time, the 5 cells (in yellow circles) were hemolysed leaving behind the cell ghosts. The scale bar is 30 *µm*.

## Conclusion

The presented method provides a practical and cost-effective solution for high-frequency acoustic microscopy by enabling the fabrication of thin-bottom petri dishes through 3D printing combined with commercially available stretch film. Our results demonstrate that this approach preserves both acoustic transparency and structural stability, making it well-suited for precise imaging applications. Beyond acoustic microscopy, the design also allows for complementary experiments, such as optical tweezing and drug response studies. This versatility enables sequential investigations of the same sample using multiple modalities. We anticipate that its usability can be further extended to other modalities, including fluorescence and confocal microscopy. Moreover, the ease of customization enables researchers to tailor the design to a wide range of experimental setups, effectively addressing the limitations of existing commercial alternatives. Overall, this technique has the potential to expand access to high-resolution acoustic imaging, particularly in resource-constrained laboratory environments.

## Ethical Statement

The experimental protocols, including blood sample, were approved by Boğaziçi University Science and Engineering Fields Human Research Ethics Committee (FMINAREK) with the ethical permission number 2025-15. Blood samples were taken using a lancet needle from the participants’ fingertips. All methods were performed in accordance with Declaration of Helsinki.

## Supplementary Materials

The design and assembly video of the custom-made petri dish can be found in [Gitlab](https://gitlab.com/tunapesen/custom-made-petri-project).

## Author Contributions

CRediT: **T.P**.: Conceptualization, Data curation, Funding acquisition, Investigation, Methodology, Project administration, Supervision, Visualization, Writing - original draft, Writing – review & editing; **B.C.E**.: Data curation, Investigation, Methodology, Visualization, Writing – review & editing; **B.A**.: Funding acquisition, Project administration, Supervision, Writing – review & editing.

## Acknowledgements

This work was made possible by the 3D printer donated to our laboratory in memory of **Barış Sadık**. We extend our heartfelt gratitude to his family for their generosity and for honoring his memory through this contribution.

## Data Availability

The data supporting the findings of this study are available on [Gitlab](http://gitlab.com/tunapesen/custom-made-petri-project).

